# No association between metabolic rate and an exaggerated sexually selected trait

**DOI:** 10.1101/2025.04.29.651352

**Authors:** Kentarou Matsumura, Tomohito Noda, Manmohan D. Sharma, Kensuke Okada, Takahisa Miyatake, David J. Hosken

## Abstract

Exaggerated sexually selected traits are thought to be energetically costly to produce and to maintain. This is especially true of sexually selected weapons, traits that are frequently positively allometric, and therefore likely to be especially energetically costly. However, there have been few studies that directly measure these putative costs. Additionally, metabolic rate, a measure of energy expenditure, is highly variable across individuals, but the causes of this variation are not always well established. Here we tested for the associations between resting metabolic rate, during development and adulthood, and the size of a weapon used during adult male-male competition in the beetle *Gnatocerus cornutus*. We also tested for genetic variation metabolic rate and weapon size. We found that metabolic rate was not statistically associated with weapon size, but there were effects of genotype and mass during development (pupal size) on weapons. Additionally, age, mass and genotype all affected metabolic rate in various ways. Despite no evidence of energetic costs to weapon development or maintenance, direct predation costs associated with them have been detected previously, and hence enlarged weapons are costly in this system.

**Significance statement:** The large weapons used in competition between males are thought to be energetically costly. However, there are rather few investigations of these putative costs. We investigate these potential costs by testing for associations between metabolic rate and weapon size in a beetle. We found no relationship between weapons and metabolism, indicating weapons are not always energetically costly.

## Introduction

Sexual selection is a major mechanism of evolution and has caused some of the most exaggerate behaviours and morphologies in nature (Andersson 1994). While sexually selected traits are typically thought to be costly, evidence for this is somewhat limited, even though understanding putative costs is important because they set limits on trait exaggeration and determine trade-offs with other fitness components (Kotiaho 2001).

Predation costs have been reasonably well documented for sexually selected characters as these traits can increase conspicuousness to predators (e.g. Endler 1987; Reznick and Endler 1982), and predators and parasitoids frequently target sexually signalling males (e.g. Ryan 1985; Hosken et al. 1984; Grey and Cade 1999; reviewed in Zuk and Kolluru 1998), even preferring the same call parameters favoured by females in some cases (Ryan 1985, 2018). Additionally, investing in sexually selected traits can generate immune function costs (e.g. Folstad and Karter 1992; Salvador et al. 1996; Hosken 2001; McKean and Nunney 2008), although it is not always clear whether reductions in elements of immune function have significant fitness impacts (Kotiaho 2001). Other physiological costs of exaggerated sexually selected traits have arguably been less studied (Halliday 1987), although energetic costs of sexual signalling have been documented (e.g. Bailey et al. 1993; Kotiaho et al. 1998; Reinhold et al. 1998), and fights can also be energetically expensive (Hack 1997; Neat et al. 1998). For example, signalling can increase metabolic rate by more than 50% (e.g. Bailey et al. 1993; Hoback and Wagner 1997), while male fighting can increase energy expenditure 6 times above resting metabolic rate (Hack 1997).

The weapons used in fighting for territories and mates are amongst the most exaggerated morphological characters, frequently scaling positively with body size, such that large individuals have disproportionately large weapons – weapons show positive static allometry (e.g. Eberhard et al. 2018; Kodric-Brown et al. 2006). This positive allometry is found in the weapons of stag beetles, earwigs and fiddler crabs for example (Kodric-Brown et al. 2006), and steep scaling relationships at least partly explains why weapons are thought to be especially costly to produce and maintain (Somjee 2021). Consistent with this, there can be considerable metabolic cost to morphological-weapon maintenance, with metabolic expenditure decreasing by about 25% when weapons are lost (Somjee et al. 2018; also see O’Brian et al. 2019). Additionally, mass-specific metabolic rates are higher in crabs with intact weapons (Allen & Levinton 2007). More generally, sexual traits are thought to be metabolically demanding (reviewed in Kotiaho 2001), and this may be especially true of long-lasting morphological characters (Allen & Levinton 2007). However, it has also been suggested that costs of weapons will depend on the tissue-types that form them (Dinh 2022; Somjee 2021). For example, rhinoceros beetle horns are hollow and therefore metabolically inactive, while fiddler crab claws are full of muscle and therefore energetically expensive (Somjee 2021). Note this relates to maintenance costs, and even when these costs are low, developmental costs, the costs of actually building large weapons, can be very significant (Rosenfeld et al. 2015). One step toward understanding the costs of sexually selected traits like weapons, is to assess the metabolic expenditure associated with them.

Metabolic rate (MR), which measures energetic expenditure on somatic maintenance (McNab 2005; Rezende et al. 2004; Snodgrass et al. 2005; Sinclair et al. 2011), could influence the resources available for investment in energetically expensive traits like weapons, with the “residual” energy not needed for basic maintenance being free to invest in these characters (Rowe and Houle 1996; Reinhold 1999; Tomkins et al. 2004; Ketola and Kotiaho 2009). Thus, MR could be negatively associated with weapon size, since lower maintenance costs could generate greater energy pools left to invest in larger weapons. Alternatively, a high MR could enable greater energy acquisition and conversion into weapons, and thus MR could be positively associated with weapon size (Bennett and Ruben 1979; Biro and Stamps 2010; Ketola and Kotiaho 2010). In either case, metabolic rate and weapon development could be linked, but the metabolic costs of weapons are highly variable (O’Brian et al. 2019), and therefore a case-by-case assessment is required. Additionally, while there is substantial within species variation in MR, explaining it has been challenging (Burton et al. 2011). This is especially true when viewed from a non-mechanistic perspective (Burton et al. 2011; Pettersen et al. 2018), and studying MR variation within an evolutionary genetics framework has been recommended (Petterson et al. 2018).

Here we tested for associations between resting metabolic rate and weapon size in the broad-horned flour beetle, *Gnatocerus cornutus*. Male beetles have enormously enlarged mandibles that are used in male-male fights (Okada and Miyatake 2009; Harano et al. 2010), and females lack this exaggerated character completely (Okada et al. 2006; Harano et al. 2010; Matsumura et al. 2024). Like many sexually-selected weapons (Kodric-Brown et al. 2006), the mandibles are positively allometric (Okada and Miyatake 2010; Shinohara et al. in review). Furthermore, males with larger mandibles have higher fighting and mating success (Okada and Miyatake 2009; Harano et al. 2010; Yamane et al. 2010; Katsuki et al. 2012), and there are also age-effects on fighting ability and a genetic component to aggression in younger males (Nishitani et al. 2024).

We have previously documented predation costs to larger mandibles, with the smaller mandibles evolving in populations subjected to male predation (Okada et al. 2021). Furthermore, this predation reduces male (but not female) longevity (Pennell et al. 2024). Thus, there are direct costs to mandibles that have life-history impacts (c.f. Kotiaho 2001), but it is not clear whether metabolic rate (MR) affects mandible size in any way. Mandible development occurs during pupation with a number of gene knockdowns just prior to pupation able to induce mandible development in females or mandible suppression in males (Sugiyama et al. 2023). We therefore tested for mandible-size/metabolic rate associations in both pupae, to assess putative energetic costs to mandible development, and adults, to test for putative maintenance costs. We did this phenotypically and using iso-female lines (Hoffmann and Parson 1989; David et al. 2005). Iso-female lines are effectively distinct but replicable genotypes, which allowed us to test for genetic effects in our traits of interest – differences between isolines indicate genetic variation in the characters that differ (Hoffmann and Parson 1989; David et al. 2005; and see e.g. Ingleby et al. 2013; Archer et al. 2024; Nishitani et al. 2024). Investigating metabolism through this lens has been recommended because genetic variation in MR may be one reason for between individual variation in this trait (Petterson et al. 2018).

Finally, we also tested for mass-MR and mass-mandible associations. There is a general expectation across species that larger animals have higher MR, but within species this may not always hold. Furthermore, if mass reflects the ability to accrue resources to spend on expensive secondary sexual traits, there could be mass-trait associations. We tested this here. While we found no associations between metabolic rate and weapon size, genotype and pupal mass influenced weapon size, and age, mass and genotype all impacted metabolic rate in various ways.

## Materials and methods

The *G. cornutus* beetle culture originated from adults collected in Miyazaki city, Japan, in 1957 (Okada et al. 2006) and has subsequently been maintained in the laboratory of National Agriculture and Food Research Organisation, Japan. The beetles used in this study were derived from that culture. They have been reared on 90% organic white flour (Doves Farm Foods) enriched with 10% brewer’s yeast (Brewers’ yeast, Thermo Fisher Scientific). Both rearing and experimentation were conducted in climate chambers maintained at 27 °C, 60% relative humidity and with a photoperiod of 14h:10h, light to dark cycle. These laboratory conditions mimic the natural conditions where *G. cornutus* was found (Katsuki et al. 2012).

In this study, we used individuals derived from 18 iso-female lines established from the stock population (Matsumura et al. 2020; Nishitani et al. 2024). To initiate these lines, 18 males and 18 females from the stock population were selected at random and paired. Subsequently, sib-sib matings within each family were used to propagate isolines for about 30 generations until the present study was conducted. As noted above, isolines represent distinct genotypes, and significant variation among them indicates genetic variation in the base population for the character of focus (Hoffmann and Parsons 1989). Previous work has shown these isolines harbour significant genetic variation for all the traits investigated even after 40 generations of propagation (e.g. Matsumura et al. 2020; Nishitani et al. 2024).

Metabolic rate (MR) and trait measurement follow standard procedures. In brief, the resting metabolic rate (RMR), assessed via CO_2_ production (see methods in Arnqvist et al. 2010; Sinclair et al. 2010; Okada et al. 2011), of know-age male pupae (N = 205: note we measured MR in many pupae and established sex on ecolsion) and adults (N = 178) was measured using flow-through respirometry (Sable Systems, Las Vegas, USA) (Lighton 2008) along with animal mass and used as our indicator of MR. Briefly, compressed zero-air (79% N_2_ and 21% O_2_) was first scrubbed of residual CO_2_ and water (Ascarite II and Magnesium Perchlorate) and regulated at a flow rate of 100ml min-1 ±1% with a Sierra Instruments (Monterey, CA, USA) model 840L mass-flow meter connected to a mass-flow control unit (Sable Systems Intelligent Mass Flow Control Unit, MFC-2, Las Vegas, USA). Air was then fed into a recently calibrated LI-COR LI-7000 infra-red gas analyzer (IRGA) capable of resolving differences of 1ppm of CO_2_ in the air, plumbed in differential mode (see Lighton 2008). Air exiting the IRGA reference cell was passed through a Sable Systems (Las Vegas USA) RM8 eight-chamber multiplexer and then forwarded to the sample cell for measurements. A PC running Sable Systems Expedata software (ver.1.8.2) and connected to a Sable Systems UI2 analog–digital interface was used for data acquisition. Within the multiplexer, one empty chamber acted as a negative control and beetles were haphazardly assigned to other chambers to minimize positional bias. Baseline measurements were made before and after each set of recordings to test and control for any drift between calibrations. Carbon dioxide production was measured for 7 minutes/beetle (with a sample read every second), and data from the most stable section (Lighton 2008) of the last two minutes of measurement (to control for wash-out) were used to calculate the average CO_2_ production for each individual (ranging from 60-100 seconds of measurement/beetle). Identifying the stable section was done manually using the ‘level’ function within Expedata software (ver.1.8.2). This function allows you to choose a contiguous set of points within the selected area of the active channel that are (for the selected number of points) most stable (flat).

After measurement of metabolic rate, pupae were placed into individually into the wells of a 24-well plate with food and maintained until they eclosed as adults, and MR was measured again at adulthood using the methods described above. To avoid an effect of feeding, adult beetles (10-15 days old: when males are sexually mature) were isolated without food for two hours before the measurement of MR. All measurements of MR were conducted between 12:00 and 19:00 in an incubator maintained at 27°C (SD±0.4_o_C) and our measure of MR (RMR) was repeatable: the intraclass correlation coefficient (ICC), estimated using the irr R package (vers. 0.84.1; Gamer et al. 2019) for 49 beetles measured thrice was 92% (ICC(C,3) = .920, 95% CI = [.872, .952] estimated using an average-score, consistency-based, two-way random effects model: F_48,96_ = 12.5; P < 0.001). This is consistent with previous repeatability tests (Matsumura et al. 2025).

Following measurement of adult MR (with MR for each individual was measured as an adult and pupae), body mass (measured as above) and mandible size were measured using standard approaches (e.g. Okada & Miyatake 2009; Harano et al. 2010; Okada et al. 2014). Briefly, male beetles were euthanized by freezing and stored at −20°C until processing for morphological measurement. Each beetle was placed on a slide and oriented consistently under a stereomicroscope (Leica M125) to ensure standardized imaging. Digital images of the whole body were captured using a mounted digital camera (Leica DFC 295) connected to a PC running ImageJ software (version 1.48). Images were calibrated to scale using a 1000μm reference standard. Mandible length, defined as the distance from the base to the tip of the mandible was quantified using ImageJ’s measurement tools. All measurements were taken by a single observer to minimize variation, and repeatability was confirmed through blind re-measurements of a subset of individuals (repeatability > 0.95, following Okada & Miyatake 2009).

### Statistics

Visual inspection of the data did not indicate significant violations of normality (and see Kneif & Forstmeier 2021; Hosken et al. 2018), and although the data-set contained one very small individual, removal of this datum did not change model outcomes (we therefore present analyses that include this male). We used two statistical approaches, one that focussed on the associations for the “populations we have (isoline as a fixed effect) and one that was more general (isoline as a random effect). For the former we used MANOVA, fixed effect GLMs, least squares linear regression and for the latter, GLMMs, with analyses were carried out in SPSS (Version 29.0.2.0 (20)) and R (R-code can be provided on request). Isoline was entered as a random effect in the GLMMs, but outcomes are unchanged if treated as a fixed effect (GLM/MANOVA). Regression was used when ignoring the effects of genotype and therefore looking at purely phenotypic associations and age was used as a fixed effect in the GLMs since our measure was very course (days). Additionally, we also calculated mass-specific MR (Reinhold 1999) for some analyses even though models that include mass and MR are recommended (Lighton 2008). Note sample sizes differ between pupal and adult analyses because not all pupae (N = 205) survived to adulthood (N = 178), and for some individuals, data were missing. Additionally, as should be the case, conclusion are unaltered when analyses are conducted with a mixed model (GLMM) approach (Supplementary Materials), so we only present the GLM/MANOVA results which are probably more intuitive.

## Results

To test for effects of MR on mandible length, we first used univariate GLMs with isoline as a fixed effect and pupal and adult mass and MR as covariates. This revealed an effect of genotype (F_17,133_ = 1.731; P = 0.044. Figure 1) and strong positive effects of pupal mass on mandible length (F_1,133_ = 8.955; P = 0.003. Figure 2), with no other predictor significantly affecting mandible length (all F < 1.63; all P > 0.2) (see Figure 3 for pupal MR and mandible size). The removal of one very small individual (Figure 1) did not alter these findings. Similarly, removal of individuals with small mandibles for their isolines (Figure 2) did not change model outcomes. Additionally, multiple regression of phenotypic values (i.e. analyses did not include “isoline”) was significant overall (F_4,150_ = 9.203; P < 0.001) because of an effect of pupal mass on mandible size (standardised *ß* = 0.315; t = 3.124; P = 0.002) with no other significant effects (adult and pupal MR, adult mass: all t < 1.3; P > 0.19). Finally, given associations between mass and MR (see below), we also ran a GLM that only included MR (adult, pupae) and isoline (fixed effect) as predictors and still found no MR-mandible associations (all F < 1.67; P > 0.19). These results are consistent with the GLMM analyses, which also showed no significant associations between metabolic rate and mandible length, but confirmed the effects of pupal mass on mandible length (see Supplementary Tables 1, 2 and 3).

**Figure 1.**
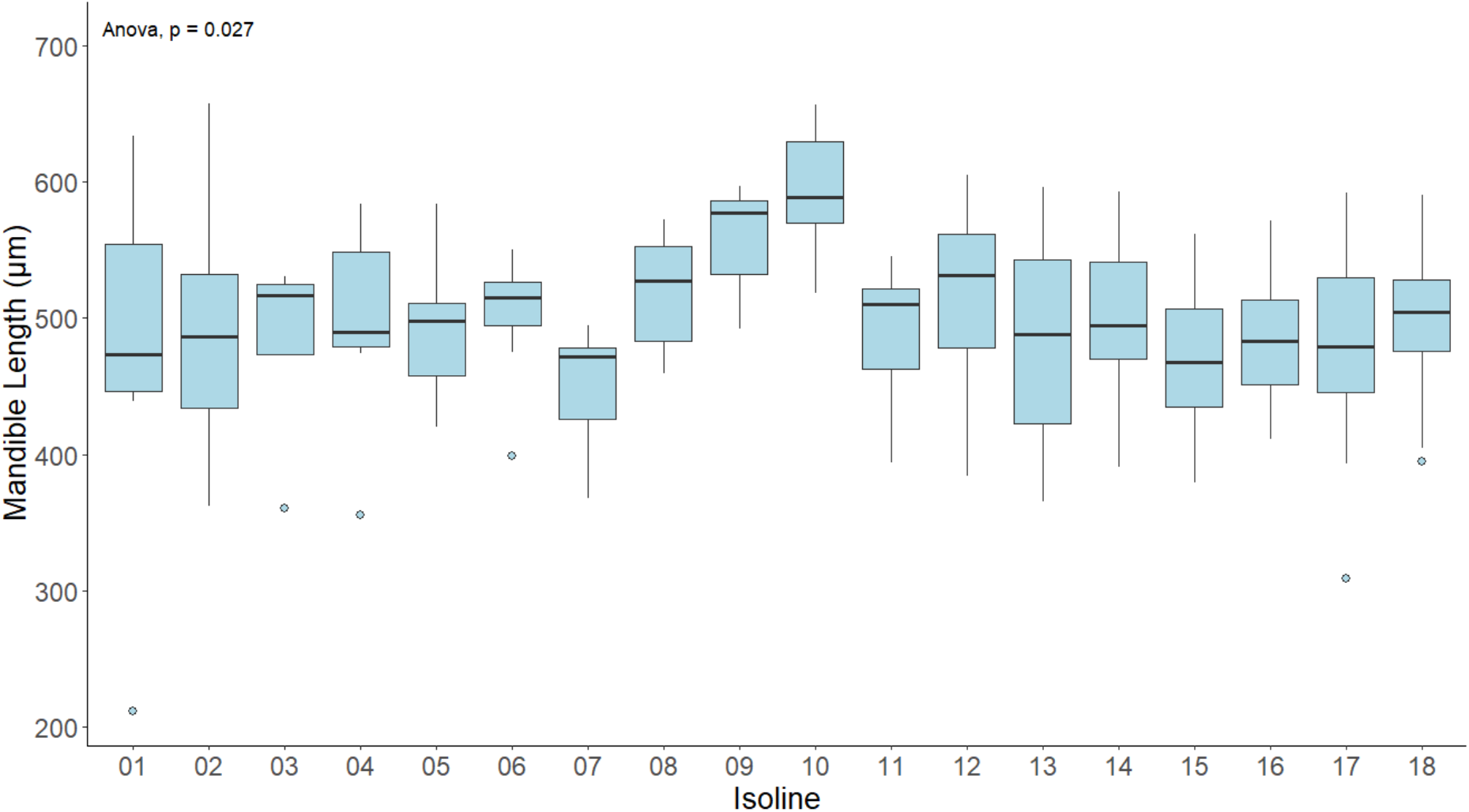
The statistically significant variation in mandible size (um) across the 18 genotypes (isoline) used in the study - genotypes have been given arbitrary numbers. Shown are medians, 1^st^ and 3rd quartile, minimum and maximum and outliers. Not ethe effect of isoline on mandible size is not changed with the removal of outliers (individuals with small mandible size for their genotype).

**Figure 2.**
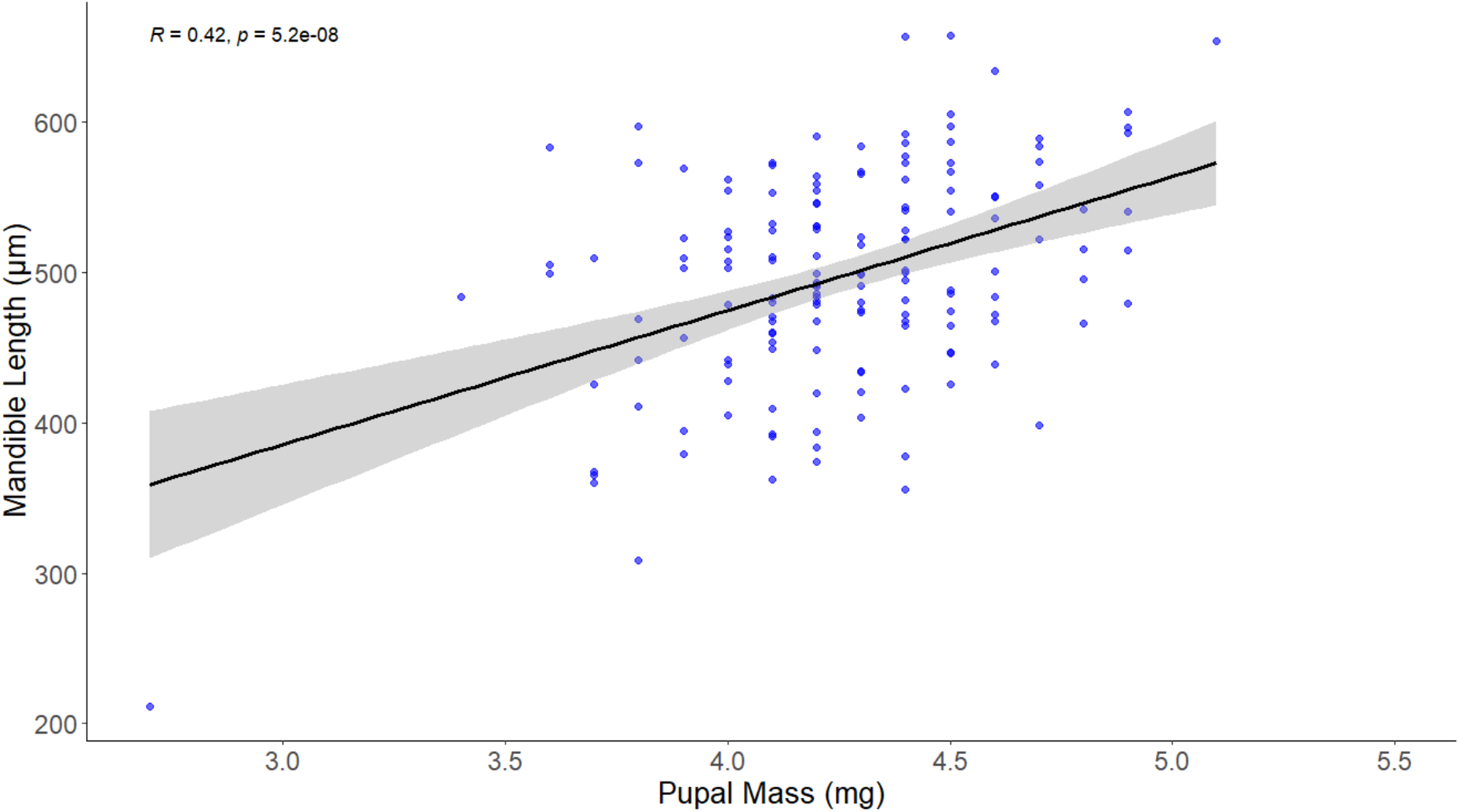
The positive association between pupal mass (mg) and the length of mandibles (µm) in adult males. This relationship is statistically significant and this is not altered by the removal of the single very small male.

**Figure 3.**
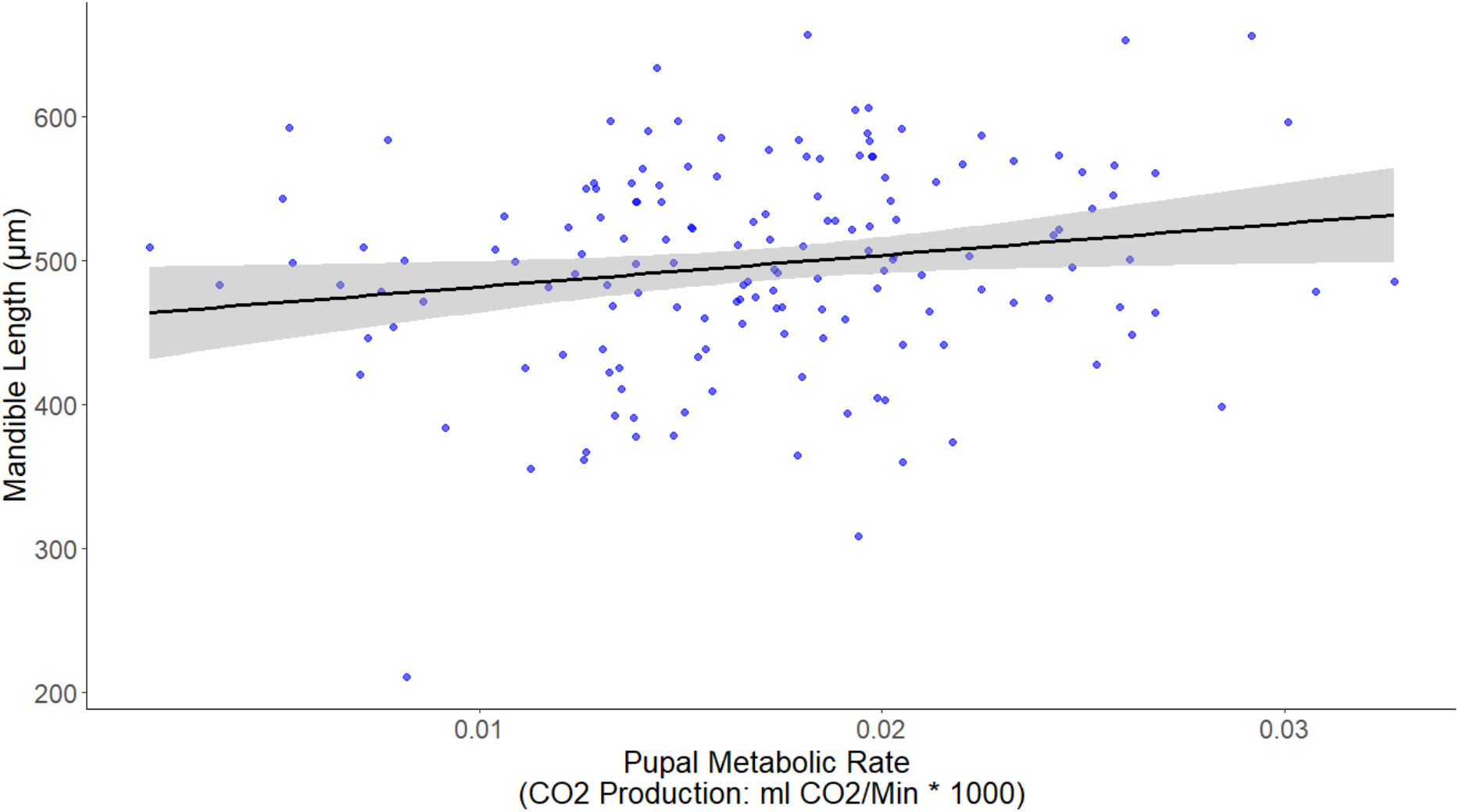
The flat non-significant association between pupal metabolic rate and mandible length. A very similar non-significant association is seen when mandible length is plotted against adult MR, and in both cases the removal of the individual with very small mandibles does not change conclusions.

We also conducted a MANOVA to test for genetic effects on the multivariate combination of our traits of interest, with genotype (isoline) as the predictor, and pupal and adult metabolic rates and mass, and adult mandible length as dependents. This showed a strong genetic effect on the multivariate combination of these traits (isoline: Wilks’ Lambda F_85,647.4_ = 1.612; P < 0.001). This was driven by genotype (isoline) effects on adult MR (F = 1.750; P = 0.041), mandible length (F = 1.857; P = 0.027) and pupal mass (F = 2.007; P = 0.015). Using mass-specific MR (Reinhold 1999) rather than mass and MR, largely did not change univariate conclusions – isoline still affects mandible length (F_17,137_ = 1.857; P = 0.027) and adult mass-specific MR (F_17,137_ = 1.84; P = 0.028). However, there was no significant effect on pupal mass-specific MR (F_17,137_ < 0.001; P > 0.999 and the multivariate effect was no longer significant (Wilks’ Lambda = 0.661; F_51,402.7_ = 1.18; P = 0.196).

We also tested for associations between mass and MR in adults and pupae using GLMs (that included isoline and age as a fixed effects) and least squares linear regression (MR vs Mass). For pupal MR, pupal age (F_3,183_ = 17.18; P < 0.001) and pupal mass (F_1,183_ = 8.50; P = 0.004) had significant effects, but there was no effect of isoline (F_17,183_ < 0.001; P > 0.99), with MR tending to increase with pupal age and mass (Figure 4). The regression of pupal MR on mass (i.e. the phenotypic association ignoring age and genetic affects) also found a significant positive association between MR and mass (standardised *ß* = 0.244; F_1,204_ = 12.84; P < 0.001). In the GLM for adult MR, adult mass did not impact adult MR (F_1,154_ = 2.092; P = 0.150), but age and genotype (isoline) had significant effects (Age: F_5,154_ = 2.698; P = 0.023. Isoline: F_17,154_ = 2.053; P = 0.011). However, the age affect was driven by the elevated MR of two 15 day-old individuals, and when they were removed from the analysis, pupal age no longer significantly impacted MR (isoline remained significant). Regression of adult MR on adult mass (i.e. the phenotypic association ignoring age and genetic affects) revealed a non-significant but positive association between MR and mass (standardised *ß* = 0.115; F_1,176_ = 2.357; P = 0.126). These results are also consistent with the GLMM analyses (see Supplementary Tables 1, 4-6).

**Figure 4.**
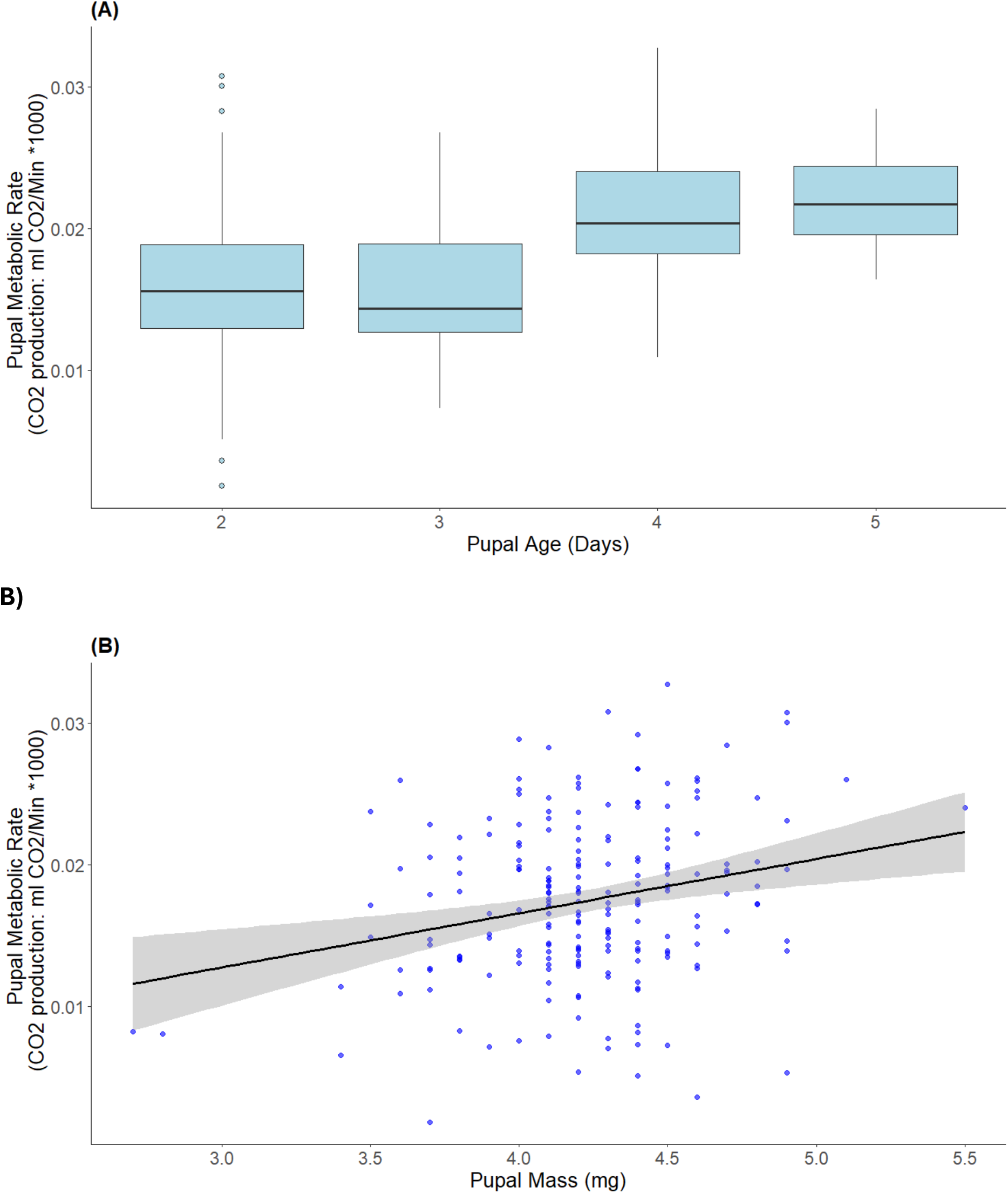
The effects of pupal age (A) and mass (B) on pupal metabolic rate (MR). MR tends to increase as development proceeds and increase as mass increases (standardised *ß* from a univariate linear regression = 0.244). In Plate A, medians, 1^st^ and 3^rd^ quartile, minimum and maximum and outliers are shown. Interpretation does not change if outliers (2-day old individuals) are removed from the analysis.

## Discussion

Despite expected associations between metabolic rate and weapon size (e.g. Tomkins et al. 2004; Biro and Stamps 2010), we did not find any statistically significant relationships between the two, either during pupation, when mandibles develop (Sugiyama et al. 2023), or during adulthood when they are used in fights (Harano et al. 2010). This is despite large sample sizes and finding a range of other statistically significant associations. Most notable of these was the significant effect of isoline on all the traits we measure, which is consistent with genetic underpinnings to these characters (David et al. 2005). We discuss these findings in turn.

There are good reasons to expect resting (basal) metabolic rate to be associated with weapons size, either because a low resting metabolic rate uses less energy and hence more energy is available to be allocated to weapons, or because a higher metabolic rate allows individuals to garner more resource to be allocated to armaments (e.g. Tomkins et al. 2004; Biro and Stamps 2010). Additionally, many weapons are positively allometric (Kodric-Brown et al. 2006), including the mandibles at the centre of our study (Okada and Miyatake 2010), and metabolic expenditure has been positively associated with weapon size in other taxa (e.g. Somjee et al. 2018). Furthermore, morphological characters are a fixed cost once developed and hence may be especially likely to be energetically expensive (Allen & Levinton 2007). However, despite all this, we did not find any associations, and MR – weapon associations are generally highly variable and appear to be influenced by the muscle mass associated with the weapons (O’Brien et al. 2019) and the tissue type that weapons are built from (Somjee 2021). In *G. cornutus*, the muscle largely used to operate the mandible sits within the pronotum and this is highly genetically and phenotypically correlated with mandible size (e.g. Harano et al. 2010). So despite not measuring muscle mass in our study, mandible size itself should capture much of the variation in muscle mass and hence reflect an MR-muscle association if it existed. However, the mandibles themselves are exoskeleton and this should reduce their metabolic maintenance costs (Dinh 2022), which is consistent with the lack of association between adult MR and weapon size in *G. cornutus*.

While there might not be significant metabolic costs associated with bearing mandibles once they have developed (Dinh 2022; Somjee 2021), we thought there could well be elevated metabolic expenditure associated with developing larger mandibles as has been found in other taxa (Somjee 2021). For this reason MR was measured during pupation – when mandibles develop (e.g. Sugiyama et al. 2023). Again however, we found no associations between (pupal) MR and weapon size, although larger pupal mass, which probably reflects resource accumulation prior to trait development, was positively associated with mandible size. If pupal mass is a reflection of resource accumulation, then it could be argued that there are energetic links to mandible size, but these are independent of pupal (and adult) metabolic rate. Consistent with this, the balance of macronutrients in the larval diet has a strong effect on mandible size in adult beetles (House et al. 2016). Additionally, in populations selected for increased mandible size, there is a reduction in survival during pupation (Okada et al. 2011), which could also mean we have underestimated metabolic expenditure on mandible growth if those individuals with the largest mandibles did not survive through pupation to be measured. In any case, predation costs associated with larger mandibles, and knock-on longevity impacts have previously been documented (Okada et al. 2021; Pennell et al. 2024). Thus there are life-history costs to larger mandibles, as required to demonstrate “real” costs associated with sexually selected characters (Kotiaho 2001).

Like all non-significant associations, it is possible that we simply lacked the statistical power to find any effect. However, our sample sizes were large, MR was repeatable, and we did find other associations, so we are reasonably confident that the lack of an association between MR and mandible size is real. This conclusion is also consistent with findings from other taxa where larger weapons are composed of more energetically inexpensive exoskeleton (Dinh 2022). However, comparing male and female mass-specific metabolic rates would provide another means of exploring potential energetic costs to mandibles (Oufiero & Garland 2007) to ensure these conclusions are robust, and this is something for future work, especially since survival through the pupal stage could be impacted by mandible size (Okada et al. 2011).

Despite finding no MR-mandible associations, we nonetheless identified isoline effects for mandible size, pupal mass and adult metabolic rate, reflecting genetic variation in all these characters (Hoffman and Parsons 1988; David et al. 2005). It was not surprising that we found evidence for a genetic basis to mandible size, since this trait has a high heritability (Okada et al. 2014), responds readily to artificial selection (Okada and Miyatake 2009) and to male specific predation (Okada et al. 2021). For example, bidirectional selection on mandible size generates rapid microevolutionary responses for both larger and smaller size, and also generates as raft of correlated response in body shape (Okada and Miyatake 2009; Harano et al. 2010). Likewise, there were apparent genetic effects on pupal mass which is not surprising as size measures are generally heritable (e.g. Messina 1993; Iyengar and Eisner 1999; Blanckenhorn and Hosken 2003). Genetic variation for MR has also been found in other taxa (e.g. Nilsson et al. 2009; Einum et al. 2019; reviewed in Pettersen 2018), although this at times depends on cyto-nuclear interactions (Arnqvist et al. 2010), but in any case, there appears to be a genetic component to adult MR in *G. cornutus*. The importance of this insight is not clear as we do not have a thorough understanding of MR-fitness links at this point. However, genetic variation in MR helps explain why MR is so variable across individuals (c.f. Burton et al. 2011). Interestingly, genotype had no effect on pupal MR, but there were strong age effects which likely reflect the effects of development stage on energy expenditure, as has been documented in other studies (e.g. Krams et al. 2017, 2018). Additionally, these age effects may have obscured any genetic effect on pupal MR. We also found no significant genotype effect on adult mass which was somewhat surprising because adult mass has a high heritability in *G. cornutus* when estimated with a paternal half-sib breeding design (Okada et al. 2014). Of course such designs are specifically to enable genetic parameter estimation and will return estimates with greater accuracy than our more general approach here.

We also documented mass-MR associations in some analyses but not others, with pupal MR having the strongest associations with mass (although MR and mass were positively associated in all analyses). While mass-MR associations are well established across species (Withers 1992; Nespolo et al. 2005), intra-specific associations can be weaker. Additionally, while some insect studies find mass-MR relationships (e.g. Lighton 2008; Okada et al. 2011; Sinclair et al. 2011; Matsumara et al. 2025), others do not (Van Voorhies et al. 2004; Canzano et al. 2006; Oikawa et al. 2006), which may reflect differences in factors like starvation resistance and time in captivity (e.g. Harshman et al. 1999; Van Voorhies et al. 2004). As noted above, age and genetic effects were also implicated in MR variation. Finally, the MR estimates we obtained here are similar to those for other similarly sized beetles (Matsumara et al. 2025).

In summary, we found no evidence for associations between metabolic rate and an exaggerated sexually selected trait used in male-male combat. This is despite the fact that this weapon shows positive allometry and has other costs associated with its expression. This finding is consistent with suggestions that tissue type affects the energetic costs of weapons (Somjee 2021). Additionally, we record genotypic effects on metabolic rate, but whether metabolic rate relates to life-history characters and fitness remains to be elucidated, as does potential associations between MR and fighting ability.

## Acknowledgements

We thank many members of FlyLab who helped in various ways with this study, especially Sarah Daisy Kendall for help with the imaging and Ruth Archer for support.

We thank Stu Bearhop and Dave Hodgson for statistical advice. DJH and MDS were funded by the Leverhulme Trust.

## Ethics

This study has ethical approval from the Ethics Committee of the Centre for Ecology and Conservation, University of Exeter.

